# Changes in coding and efficiency through modular modifications to a One Pot PURE system for in vitro transcription & translation

**DOI:** 10.1101/2023.07.28.550900

**Authors:** Phuoc H. T. Ngo, Satoshi Ishida, Bianca B. Busogi, Hannah Do, Maximiliano A. Ledesma, Shaunak Kar, Andrew Ellington

## Abstract

The incorporation of unnatural amino acids are attractive methods for improving or bringing new and novel functions in peptides and proteins. Cell-free protein synthesis using the Protein Synthesis Using Recombinant Elements (PURE) system is an attractive platform for efficient unnatural amino acid incorporation. In this work, we further adapted and modified the One Pot PURE for a robust and modular system of enzymatic single site-specific of unnatural amino acid. We demonstrated the flexibility of this system through the introduction of two orthogonal aminoacyl tRNA synthetases and the suppression of two distinctive stop codons.

## INTRODUCTION

The incorporation of non-canonical amino acids into proteins has greatly expanded their functionalities,^1,2^ including new-to-nature catalytic functions,^3–5^ enhancement of preexisting properties,^6,7^ and probing of enzymatic activities.^8,9^ To this end, cell-free protein synthesis (CFPS) has emerged as an attractive means of readily introducing non-canonical amino acids. By eliminating membrane permeability as a barrier to substrates and by reducing competition with endogenous machineries, it has proven possible to greatly expand genetic code modifications.^2,10,11^ In particular, the Protein synthesis Using Recombinant Elements (PURE) system relies on reconstituting individually purified transcription and translational factors,^2,11–13^ and allows the ready programming of protein production through the simple introduction of template DNA. Relative to lysate-based methods, the PURE system contains less protease and nucleases activity, which greatly enhances translation, and leads to less batch-to-batch variation.^11^

Unfortunately, the PURE system suffers from high cost and / or requires intensive efforts to produce in a laboratory setting.^14^ Maerkl and coworkers have addressed these issues by developing a robust and low-cost method, termed One Pot PURE, in which individual overexpression strains for each of 36 translation and transcription factors are generated during a single co-culture, and purified jointly via a nickel chelate column,^14^ greatly reducing the labor, cost, and time for producing proteins in vitro at even higher yields than the canonical PURE system. Because of the inherent modularity of introducing translation factors, it should be possible to adapt the One Pot PURE system for the incorporation of non-canonical amino acids through the simple expedient of introducing an engineered orthogonal aminoacyl tRNA synthetase into the coculture. Here, we demonstrate the basal incorporation and optimization of translation for two non-canonical amino acids (ncAA), levodopa (L-DOPA) and N-ε-propagyloxycarbonyl-L-lysine (ProgK) via amber and ochre stop codons, respectively.

## RESULTS AND DISCUSSION

### Modifying the One Pot PURE system for more stable expression

Initial attempts to utilize the One Pot PURE system revealed batch-to-batch variability that could be traced to the mutation of T5 promoter elements driving the expression of transcription and translation factors (**Figure S1**). Among the 16 components that utilized either the pQE30 or pQE60 backbones, a T5 promoter deletion (around the -10 region) was found in 12 plasmids following two weeks of storage as frozen glycerol cell stocks and subsequent outgrowth. This issue could in turn be ascribed to the fact that the pQE30 and pQE60 series plasmids, upon which much of the system was built, do not themselves encode genetic repression of translation factor expression, and thus loss or variation in copy number of a second, LacI-expressing plasmid (pREP4) may lead to undue metabolic stress on at least some of the strains during the growth of the system as a whole. This issue has also been noted by Pluckthun and Jia, separately^15,16^.

To overcome this deficiency, we attempted to more stably repress expression of transcription and translation factors. Two strategies were explored. First, we expressed all genes under the control of the T7 RNA polymerase promoter and an adjacent TetR operator and included the Tet repressor on each expression plasmid (the so-called pKAR2 series of vectors^17^, **Figure 1a**). The pKAR2 plasmids were transformed into a B21(DE3) strain for expression; an additional advantage of this configuration is that the T7 RNA polymerase promoter in the genome is under the control of LacI, allowing for two levels of control for expression of any individual translation factor (via IPTG and tetracycline induction). Second, the original pQE30 and pQE60 plasmids encoding translation factors were transformed into an *E. coli* BL21 Marionette strain, which contains a genome-encoded array of 12 different, constitutively expressed repressors including LacI^18^ (**Figure 1a**). These reconfigured systems were compared with a pristine version of the original One Pot PURE system (M15).

**Figure 1.**
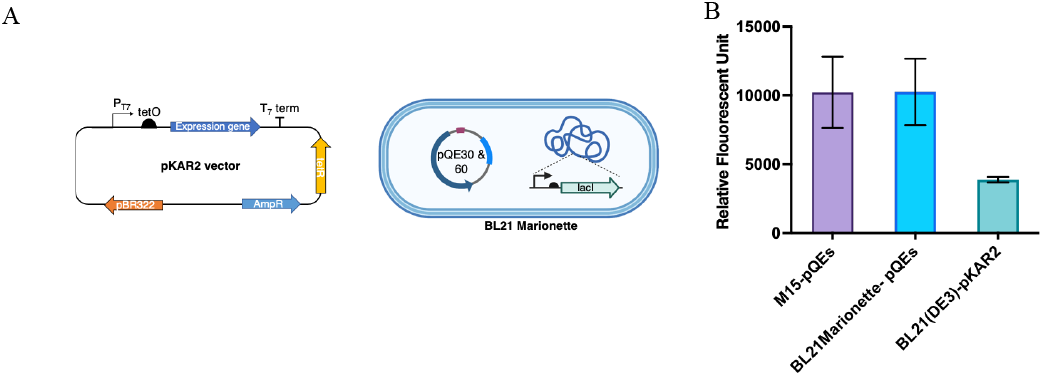
(A) Alternative protein expression strategies included changing into a more repressible plasmids (pKAR2-left) and changing into more compatible E. coli strain with genetically encoded repressors (BL21 Marionette-right). (B) the activity of purified protein factor was assessed via in vitro transcription and translation of sfGFP reporter upon reconstitution

For each system, the 36 strains were grown together, copurified translation factors, and tested for *in vitro* transcription and translation (IVTT) assays with independently purified ribosomes and using sfGFP as a reporter. Expression in the BL21 Marionette strain yielded an IVTT mixture as active as the original mixture, while T7 RNAP-based expression yielded a less active mixture, at least under the induction conditions utilized (**Figure 1b)**.

### Non-canonical amino acid incorporation in One Pot PURE via amber codon suppression

One potential utility of the One Pot PURE system would be for rapidly testing the abilities of orthogonal tRNA synthetase (aaRS):tRNA pairs for augmenting the genetic code. Previously, our lab has developed an orthogonal aaRS:tRNA pair capable of incorporating L-DOPA (**Figure 2b**) via Amber codon suppression^19^, and a corresponding reporter for L-DOPA incorporation, an orange fluorescent protein (OFP) derivative (**Figure 2a)**^19^. The OFP reporter contains mutations at S205C and an in-frame Amber stop codon at the Y66 position, and only successful incorporation of L-DOPA yields a signal that is red-shifted in both excitation and emission maxima wavelength (ex/em: 518/526 nm) comparing to the wildtype sfGFP (ex/em: 490/508 nm). Without L-DOPA, no signal would be observed at the expected excitation and emission wavelength of OFP nor sfGFP. A L-DOPA aaRS with a C-terminal histidine tag expression construct (via a BL21 (DE3) strain and the pKAR2 vector containing an inducible T7 RNAP promoter) was introduced into a One Pot culture, while the strain producing RF-1 (Release Factor-1, that normally mediates termination at Amber codons) was excluded. The suppressor tRNA was prepared in parallel by *in vitro* transcription of a linear DNA template and purification via denaturing gel electrophoresis.

**Figure 2:**
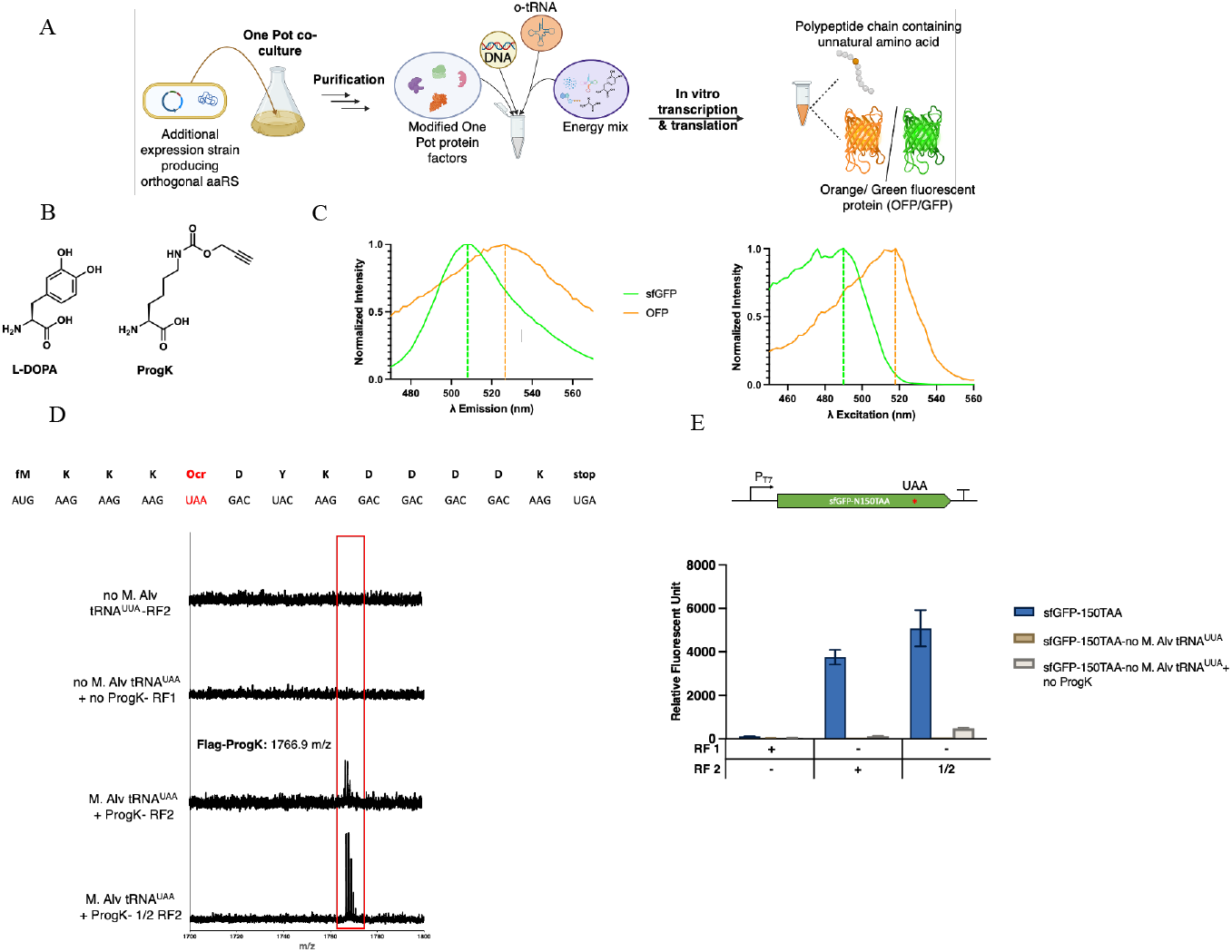
(A) Overall schematic of producing customized One Pot protein factors for in vitro transcription and translation to synthesize wild-type GFP, or to incorporate unnatural amino acid into short peptide chain, or into orange fluorescent protein depending on the input of different plasmid. (B) Unnatural amino acids used in this study, levodopa (L-DOPA) and N-χ-propagyloxycarbonyl-L-lysine (ProgK). (C) Emission (left) and excitation (right) spectrum of the IVTT reaction indicating red-shift in λ_max_of orange fluorescent protein (orange line) comparing to sfGFP (green line). (D) Peptide sequence containing in-frame ochre codon (UAA) and N-terminal Flag epitope (top); MALDI-MS spectrum (bottom) of ochre codon suppression for ProgK incorporation using modified One Pot protein factors (ΔRF1, +RF2); e) Schematic of sfGFP-N150TAA reporter for ProgK incorporation (top) and fluorescent data (ex/em: 480/512 nm) taken from IVTT reaction using modified One Pot protein factors (ΔRF1, +RF2).

The reconstituted system that included both the orthogonal aaRS and purified suppressor tRNA was prepared, and the *in vitro* transcription and translation of a small C-terminal FLAG-tagged peptide containing an in-frame amber codon was initially carried out. MALDI-TOF analysis indicated successful incorporation of L-DOPA with high fidelity (no tyrosine misincorporation was observed; **Figure S2**). Plasmids encoding either the OFP reporter with an Amber codon or a wild-type GFP reporter (Supplementary Information) were also introduced into the reconstituted IVTT reaction that contained the DOPA aaRS. The excitation and emission spectrum following translation indicated a red shift in λ_max_for the Amber-suppressed, DOPA-containing OFP compared to the wild-type GFP, indicating the successful incorporation of DOPA into the protein (**Figure 2c**).

### Capturing the next stop codon for efficient noncanonical amino acid incorporation

To demonstrate the adaptability and programmability of the system, we attempted to suppress the other two stop codons, opal (UGA) and ochre (UAA). This expansion would also potentially allow us to enzymatically incorporate two distinct non-canonical amino acids using the One Pot PURE system. We first examined the suppression of the opal stop codon using an orthogonal pyrrolysyl aminoacyl-tRNA synthetase:tRNA pair^20,21^. Since the opal codon (UGA) is only recognized by the Release factor-2 (RF-2),^22^ we eliminated the strain expressing RF-2 from the One Pot co-culture to prevent termination that would give a truncated protein, while including the strain expressing the orthogonal pyrrolysyl aminoacyl-tRNA synthetase (PylRS). This modular ability to delete and include factors at will is a major advantage of the One Pot system.

We chose to test orthogonal aaRS variants from *Methanosacrina Barkeri* (*M. Barkeri*) and *Methanomethylophilus Alvus* (*M. Alvus*) for the incorporation of N-χpropagyloxycarbonyl-L-lysine (ProgK) (**Figure 2b**) and evaluated *in vitro* transcription and translation and suppression of an in-frame stop codon in a short peptide containing a C-terminal epitope FLAG tag (**Figure S3**). The M. Barkeri PylRS was chosen as it had previously been shown to accept a wide variety of non-canonical amino acids,^20^ while the *M. Alvus* PylRS has been shown to be more tractable for engineering, as it lacks an N-terminal domain ^21,23–25^ that leads to solubility issues for the *M. Barkeri* PylRS^20,26^. Only the *M. Alvus* PylRS yielded the desired peptide product in the One Pot system (**Figure S3**).

However, there was noticeable misincorporation of tryptophan across from the opal codon, even with the addition of purified orthogonal suppressor tRNA^UCA^ and ProgK (**Figure S3**). Tryptophan misincorporation was likely due to wobble base-pairing, given the similarity of the tryptophan (UGG) and opal (UGA) codons, and had been observed in previous reports.^27–29^ Thus, to potentially increase discrimination, we changed the suppression codon to be an ochre stop codon (UAA) and also changed the anticodon of the orthogonal *M. Alvus* Pyl tRNA to respectively UUA (*M. Alv* tRNA^UUA^). To demonstrate ProgK incorporation we used a sfGFP reporter that contained an in-frame ochre stop codon at the 150th amino acid and a final opal termination codon (sfGFP-N150TAA). Ochre codons are suppressed by both RF1 and RF2 in roughly equal measure. ^30,31^ To improve suppression we eliminated RF1, but retained RF 2 to ensure proper termination and help eliminate background incorporation of canonical amino acid.^22^ A FLAG peptide product containing ProgK and no misincorporation of natural amino acids was observed (**Figure 2d**). We attempted to further improve suppression, while retaining fidelity, by lowering the amount of RF 2,^22^ which again could be readily accomplished in the One Pot system by simply decreasing the volume of cells expressing RF2As a result. One Pot mixtures containing ½ volume of RF yielded higher amounts of active sfGFP protein with only a slight increase in background (**Figure 2e**). Further dilution of RF2 did not significantly improve background (**Figure S4**) nor yield (**Figure S5**).

Overall, we adapted the One Pot PURE system for the incorporation of non-canonical amino acids and demonstrated the suppression of both an amber (UAG) and ochre (UAA) stop codon in a single mRNA. The components of the One Pot system are easily customizable in a way that the PURE system on its own is not ^15,16^; different factors can be readily introduced and titrated through the simple expedient of changing the relative amounts of inoculants in the co-culture.

## METHODS

### Materials

All chemicals and biologicals reagents were purchased without any further purification. LB medium was purchased from Fisher Sci. Carbenicillin antibiotic and Isopropyl β-D-1-thiogalactopyranoside (IPTG) were purchased from Golden Bio. Levodopa (L-DOPA), N-ε-propargyloxycarbonyl-L-lysine (ProgK), and anhydrotetracycline (aTC) were purchased from Sigma-Aldrich. Ni-NTA resin, dialysis cassettes (3.5K MWCO), and gravity flow column were purchased from Thermofisher. Amicon Ultracentrifugal Filters (3K MWCO), Anti-Flag M2 agarose, ZipTip desalting pipette tip, and E. Coli tRNA were purchased from Sigma-Aldrich. All buffers used are listed in Table S2 and S3. 2-Mercaptoethanol was added immediately before use.

### Escherichia coli Strains, Plasmids, linear DNA template, and tRNA

E. Coli BL21(DE3) strain (purchased from New England Biolab, NEB), BL21Marioentte (purchased from Addgene, cat. 108253), and M15 (generously donated from Professor Michael Jewett’s research group) were used for protein expression. For cloning and propagating plasmids, Dh10bMarionette (purchased from Addgene, cat. 108251) was used to maintain the integrity of the pQE-series plasmid.

Plasmid pJL1-sfGFP was purchased from Addgene (cat. 69496). All the expression plasmids to construct One Pot protein factors were purchased through Addgene (cat. 124103-124138). Plasmids encoding M. Barkeri PyLRS and M. Alv PylRS pET_BCS_barkeri and pET-AlvusPylRS-CtermHIS were generously donated by Professor Michael Jewett’s research group. pKAR2-Mj.DOPARs was cloned using pKAR2DO dropout plasmid and synthetic DNA gene blocks or gel-purified PCR product from amplifying pQEs and pETs series plasmids containing appropriate BsaI cut-site and overhang for Golden Gate cloning.

The DNA template used for *in vitro* transcription was prepared by overlap PCR extension from primers in table S5. The PCR product was precipitated using phenol:chloroform:isoamyl alcohol extraction and ethanol precipitation, suspended with nuclease-free water and directly used with HiScribe T7 RNA Polymerase kit (purchased from NEB). Full-length of transcribed product was purified via TBE:Urea PAGE gel, precipitated with ethanol, and resuspended with nuclease-free water. Quantification of tRNA was used via NanoDrop. DNA template encoding short peptide containing in-frame stop codon and FLAG epitote was prepapred with overlap PCR extension from primers in table. The PCR product was precipitated using phenol:chloroform:isoamyl alcohol extraction and ethanol precipitation, suspended with nucleasefree water and directly used in CFPS.

### Preparation of His-tag gravity column

Ni-NTA resin (2 mL) is added into gravity flow chromatography column. The resin was then equilibrated with buffer A (17.5 mL) and ready to use.

### Modified preparation for OnePot protein purification

Preparation for OnePot protein purification in this work was adapted from previous report^14^ and slightly modified. Cultures for protein expression were all inoculated in LB broth, supplement with 100 μg/mL carbenicillin and 50 μg/mL Kanamycin (only when M15 strain was used), and grown at 37°C, 240 rpm. All overnight and saturated cultures were inoculated in 0.3 mL of LB in a 96 deep-well plates. The strain expression EF-Tu and orthogonal aaRS for unnatural amino acid incorporation were inoculated and grown in 3 mL of LB in a standard 14 mL culture tube. The composition of One Pot expression culture at 250 mL of LB media in 500 mL flask followed Table S1. The expression culture was first grown for 2h before induction with 0.1 mM IPTG and 100 ng/mL (only when pKAR2 series plasmid was added in One Pot expression culture) for 3h. Cells were then harvested by centrifugation (4000 xg, 10 minutes, 4°C) and stored at -80°C overnight.

Frozen cells were allowed to thaw on ice for 5 minutes from -80°C and resuspended in 10 mL of Buffer A. The suspended cell pellets were transferred into a 50 mL Falcon tube. Cells were lysed using sonication method on ice with addition of splashes of brine (10s ON/ 10s OFF, 35% amplitude, ∼2000 kJ delivered or 2 minute and 40 seconds). Lysed mixture was then transferred to Nalgene Oak Ridge centrifuge tube. Cell debris was removed by centrifugation (15,000 rpm, 20 minutes, 4°C). Supernatant was then transferred to a prepared His-tag (as described above) gravity column and incubated for 3 hours in 4°C with constant shaking. After incubation, unbound lysate was allowed to flow through the column. It was then washed with 25 mL of Wash Buffer (95% buffer A, 5% buffer B) and eluted with 3 mL of Elution Buffer (10% buffer A, 90% buffer B). The purified lysate was then dialyzed using 3500 MWCO (Molecular Weight Cut Off) dialysis cassette overnight in HT buffer at 4°C.

In the final day of purification, dialyzed solution was added into 15 mL Amicon Ultra filter with 12.5 mL of HT buffer for the final buffer exchange. The solution was concentrated to ∼1 mL using centrifuge at 4000 rpm in 4°C. The protein solution was mixed with Storage Buffer B in equivalent amount. Another buffer exchange was accomplished using 0.5 mL Amicon Ultracentrifugal filter unit at 14,000 rpm in 4°C for 45 minutes. Purified protein mixture was collected and quantified using Bradford Assay with bovine gamma globulin as standard (detailed description in the subsequent section). Working protein factor was diluted or concentrated to 12,250 ng/mL and stored in -80°C for further usage.

### Ribosome purification

E. Coli A19 strain was grown overnight in 100 mL of LB broth overnight at 37°C, 270 rpm. On the next day, 30 mL of overnight culture inoculated in 2L of LB media. The culture was grown at 37°C, 270 rpm until the O.D 600= 0.6-0.8. Cells were collected via centrifuge (4000 RCF, 20 minutes, 4°C) and weighed. If the combined mass of the cell pellets is below 10 grams, it is necessary to grow more culture until the cell weighs more than 10 gram. After harvesting, cell pellets were then stored at -80°C until the next day. Frozen cell pellets were lightly thawed on ice and resuspended in 50 mL of suspension buffer and transferred to a Nalgene Oak Ridge centrifuge tube. Cell was then lysed by sonication (10s ON/ 10s OFF, 40% amplitude, 4 minutes) on ice with additions of splashes of brine. Cell debris was then removed via centrifugation (20,000 RCF, 20 minutes, 4°C). Collected supernatant was fractionated by the addition of equivalent amount of high salt buffer. Lysate was clarified one more time via centrifugation (20,000 RCF, 20 minutes, 4°C) and filtered through 0.45 μm PTFE filter.

Subsequently, ribosome was purified using hydrophobic chromatography (3×5 mL HiTrap Butyl HP column (VWR)) with Fast Performance Liquid Chromatography (FPLC) (Akta Purifier) at 4°C and with a flow rate of 4 mL per minute. Prepared lysate was loaded onto the column after equilibrating with 60 mL of buffer C. Purification follows with washing with 45 mL of buffer 1 (100% buffer C) followed by 75 mL of wash buffer 2 (80% buffer C, 20% buffer D). Elution of ribosome with 60 mL of ribosome elution buffer (50% buffer C, 50% buffer D) following by 60 mL of final elution buffer (100% buffer D). Fractions of ribosome contained those with strong absorbance at 280 nm and pooled together. The column can be recovered by washing with 1 M NaOH. and 0.1 M acetic acid, then stored in 20% ethanol.

With ultra-centrifugal polycarbonate tubes of 32 mL void volume, 15 mL of cushion buffer was added first. Recovered fraction of ribosome was then overlaid on top of the cushion buffer with equivalent amount of cushion buffer. The ribosome was pelleted by ultracentrifugation (100,000 RCF, 16h, 4°C). The pellets were then washed two times with 500 uL of ribosome buffer and resuspended with a magnetic stirrer with 200 uL of ribosome buffer. The ribosome solution was concentrated using Amicon Ultra filter unit with 3000 MWCO by centrifugation (14,000 RCF, 10 minutes, at 4°C). The concentration of ribosome was measured using absorbance at 260 nm. For an absorbance of 10 with 1:100 dilution, the concentration corresponds to 23 uM. The final ribosome working solution was diluted to 10 uM and stored in -80°C for further usage.

### Bradford Assay for Protein Quantification

Bovine gamma globulin was used as a standard. Calibration curve was constructed ranging from 0 to 20 ug/mL. For measurement, purified protein mixture was diluted 1:15, and 5 uL of diluted sample was added into 250 uL of Bradford reagent in 96-well microassay plate. Plate was then incubated at room temperature with light shaking to avoid formation of bubbles. Absorbance at 595 nm was then measure using plate reader.

### Cell-free protein synthesis and characterization

For 10 μL reaction scale without unnatural amino acid incorporation, μL of 4x Energy mix (Table S4) was mixed with 1.8 μL of 10 μM Ribosome and 1.3 μL of One Pot protein factors. DNA plasmid template was added with the final concentration of 50 ng/uL. For unnatural amino acids incorporation, purified orthogonal tRNA (o-tRNA) and unnatural amino acid were added for a final concentration of 5 μM and 0.25 mM respectively. The mixture was brought to a final volume of 10 μL with addition of nuclease and protease-free water.

Short peptide containing C-terminal Flag epitote was purified via anti-FLAG agarose resin from the CFPS reaction mixture. The purified peptide was then de-salted and subjected to MALD-MS for analysis.

Quantificaiton of sfGFP expression level were carried out by incubating the CFPS reaction at 37°C in a plate reader (Infinite 200 Pro, Tecan) for 3h and measured (excitation and emission: 488 nm and 512 nm for sfGFP). For taking emission (fixed excitation wavelength at 425 nm, and emission scan from 460 to 630 nm) and excitation scan (fixed emission wavelength of 600 nm, and emission scan from 460 to 560 nm) of OFP, the CFPS reaction was incubated 37°C in a plate reader for 5h and measured.

### Materials Availability

All plasmids in this study are available on Addgene.

## ASSOCIATED CONTENT

### Supporting Information

The Supporting Information is available free of charge on the ACS Publications website.

Sequence for observed T5 promoter deletion; MALDITOF MS spectrum of the translated peptide containing levodopa unnatural amino acid; MALDI TOF-MS spectrum of purified Flag peptide containing an in-frame opal codon from IVTT reactions using One Pot protein factor (ΔRF 2, +M. Alvus Pyrrolysyl aaRS); MALDI-MS spectrum of purified Flag peptide containing an in-frame opal codon from IVTT reactions using One Pot protein factor (ΔRF 1, +M. Alvus aaRS); Cell-free protein synthesis of N-ε-propagyloxycarbonyl-L-lysine containing sfGFP using modified One Pot protein factors containing ¼, 1/8, and 1/16 volume of RF 2; Table for olume of each cells expressing various transcriptional and translational enzymes to be added into the One Pot co-culture for 250 mL scale; Buffer recipe for One Pot protein factor purification; Buffer recipe for ribosome purification; Energy mixture recipe; Primers sequence to generate DNA template for tRNA and short peptide containing C-terminal FLAG epitote; Protein sequences of each reporters used in this study. (PDF)

## Notes

The authors declare no competing financial interest.

## ACKNOWLEDGMENT

This research was sponsored by grants from the Army Research Office (W911NF-16-1-0372 P00015, FABSUBW911NF23P0004) and a grant from The Welch Foundation (F-1654). This research was also sponsored by the Army Research Office and was accomplished under Cooperative Agreement Number W911NF-22-2-0246. The views and conclusions contained in this document are those of the authors and should not be interpreted as representing the official policies, either expressed or implied, of the Army Research Office or the U.S. Government. The U.S. Government is authorized to reproduce and distribute reprints for Government purposes notwithstanding any copyright notation herein.

## ABBREVIATIONS

PURE: Protein Synthesis Using Recombinant Elemtns
CFPS: Cell-free protein synthesis
IVTT: In vitro transcription and translation
aaRS: aminoacyl-tRNA synthetease
ncAA: noncanonical amino acids
L-DOPA: levodopa
ProgK: and N-ε-propagyloxycarbonyl-L-lysine
OFP: orange fluorescent protein
PylRS: pyrrolysyl-tRNA synthetase
RF-1: Release Factor-1
RF-2: Release Factor -2
M. Barkeri: Methanosacrina Barkeri
M. Alvus: Methanomethylophilus Alvus
sfGFP: superfolder green fluorescent protein

## For tables of content only

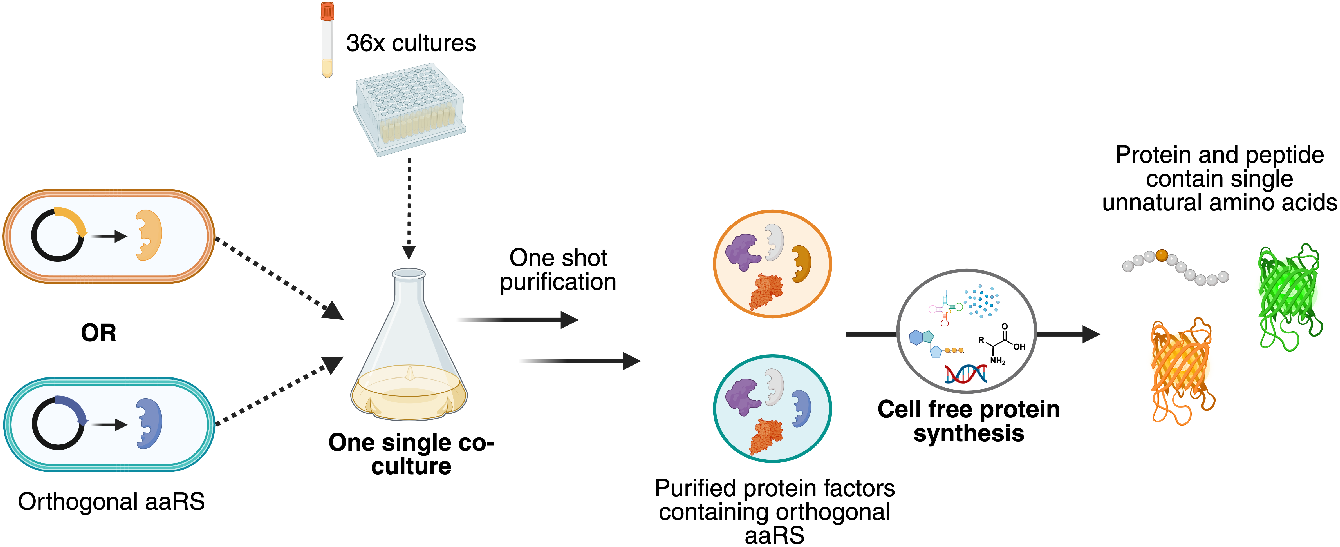

