## Supplementary Information for "Changes in coding and efficiency through modular modifications to a One Pot PURE system for in vitro transcription & translation"

| | |
|---|---|
| **Intact T5 promoter/operator** | TTGTGAGCGGATAACAATTATAATAGATTCAATTGTGAGCGGATAACAATTTCACACAGAATTCATT |
| **Sequencing result** | TTGTGAGCGGATAACAATT-----------------------------------------------------------TCACACAGAATTCATT |

**Figure S1:** T5 promoter deletion observed in M15 strains with pQE30/60 plasmid system.

| **fM** | **K** | **K** | **K** | **Amb** | **D** | **Y** | **K** | **D** | **D** | **D** | **D** | **K** | **stop** |
|---|---|---|---|---|---|---|---|---|---|---|---|---|---|
| AUG | AAG | AAG | AAG | UAG | GAC | UAC | AAG | GAC | GAC | GAC | GAC | AAG | UGA |

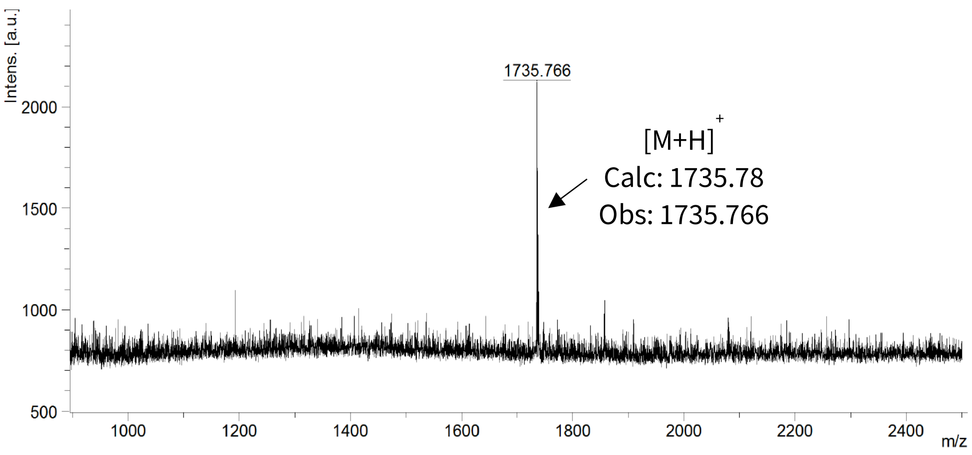


**Figure S2:** MALDI-TOF MS spectrum of the translated peptide containing levodopa unnatural amino acid.

| fM | K | K | K | Opl | D | Y | K | D | D | D | D | K | stop |
|---|---|---|---|---|---|---|---|---|---|---|---|---|---|
| AUG | AAG | AAG | AAG | UGA | GAC | UAC | AAG | GAC | GAC | GAC | GAC | AAG | UAG |

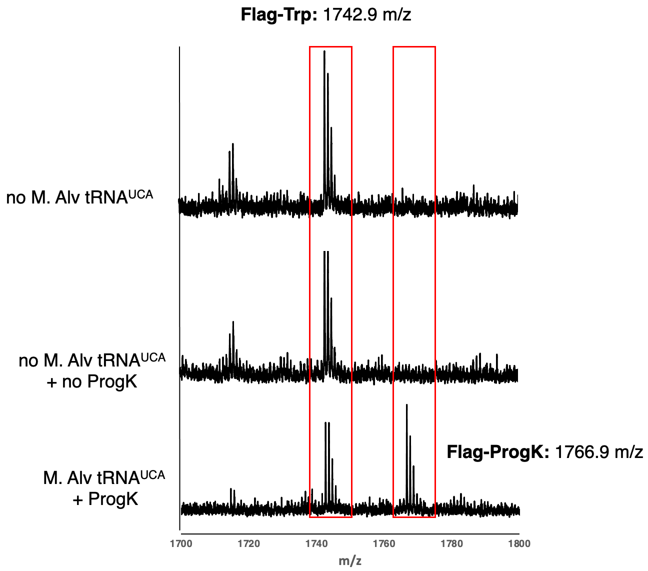


**Figure S3**: MALDI TOF-MS spectrum of purified Flag peptide containing an in-frame opal codon from IVTT reactions using One Pot protein factor (ΔRF 2, +M. Alvus Pyrrolysyl aaRS). Calculated m/z of Flag-Trp: 1742.81 and Flag-ProgK: 1766.83

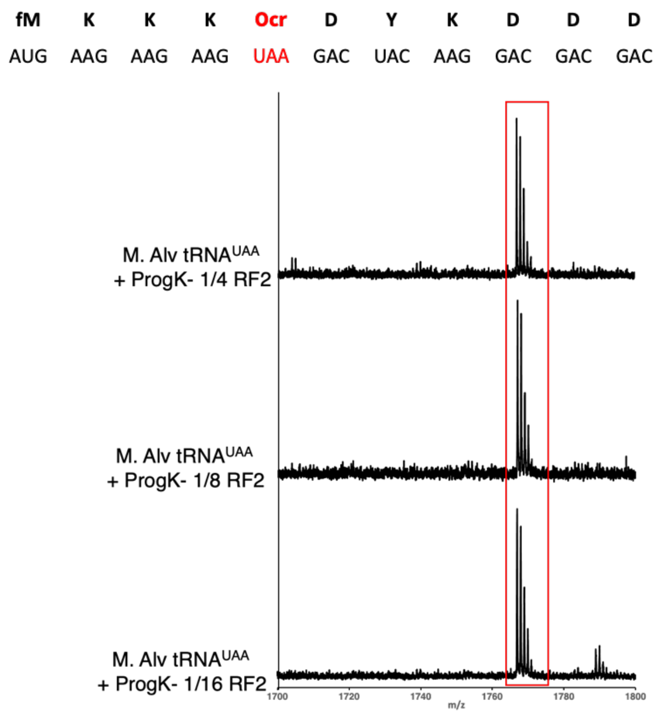


**Figure S4:** MALDI-MS spectrum of purified Flag peptide containing an in-frame opal codon from IVTT reactions using One Pot protein factor (ΔRF 1, +M. Alvus aaRS). Calculated m/z of Flag-ProgK: 1766.83.

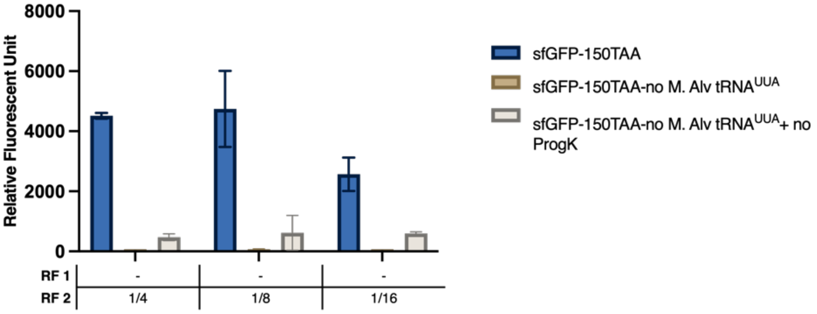


**Figure S5:** Cell-free protein synthesis of N-ε-propagyloxycarbonyl-L-lysine containing sfGFP using modified One Pot protein factors containing ¼, 1/8, and 1/16 volume of RF 2.

**Table S1:** Volume of each cells expressing various transcriptional and translational enzymes to be added into the One Pot co-culture for 250 mL scale. All volume indicated above are in µL units.

| **Sample Name** | **Standard** | **ΔRF 1+ orthogonal aaRS** | **ΔRF 2+ orthogonal aaRS** | **1/2 RF 2 + orthogonal aaRS + ΔRF 1** | **1/4 RF 2 + orthogonal aaRS + ΔRF 1** | **1/8 RF 2 + orthogonal aaRS+ ΔRF 1** | **1/16 RF 2 + orthogonal aaRS + ΔRF 1** |
|---|---|---|---|---|---|---|---|
| AlaRS | 27.5 | 27.5 | 27.5 | 27.5 | 27.5 | 27.5 | 27.5 |
| ArgRS | 27.5 | 27.5 | 27.5 | 27.5 | 27.5 | 27.5 | 27.5 |
| AsnRS | 27.5 | 27.5 | 27.5 | 27.5 | 27.5 | 27.5 | 27.5 |
| AspRS | 27.5 | 27.5 | 27.5 | 27.5 | 27.5 | 27.5 | 27.5 |
| CysRS | 27.5 | 27.5 | 27.5 | 27.5 | 27.5 | 27.5 | 27.5 |
| GlnRS | 27.5 | 27.5 | 27.5 | 27.5 | 27.5 | 27.5 | 27.5 |
| GluRS | 27.5 | 27.5 | 27.5 | 27.5 | 27.5 | 27.5 | 27.5 |
| GlyRS | 27.5 | 27.5 | 27.5 | 27.5 | 27.5 | 27.5 | 27.5 |
| HisRS | 27.5 | 27.5 | 27.5 | 27.5 | 27.5 | 27.5 | 27.5 |
| IleRS | 27.5 | 27.5 | 27.5 | 27.5 | 27.5 | 27.5 | 27.5 |
| LeuRS | 27.5 | 27.5 | 27.5 | 27.5 | 27.5 | 27.5 | 27.5 |
| LysRS | 27.5 | 27.5 | 27.5 | 27.5 | 27.5 | 27.5 | 27.5 |
| MetRS | 27.5 | 27.5 | 27.5 | 27.5 | 27.5 | 27.5 | 27.5 |
| PheRS | 27.5 | 27.5 | 27.5 | 27.5 | 27.5 | 27.5 | 27.5 |
| ProRS | 27.5 | 27.5 | 27.5 | 27.5 | 27.5 | 27.5 | 27.5 |
| SerRS | 27.5 | 27.5 | 27.5 | 27.5 | 27.5 | 27.5 | 27.5 |
| ThrRS | 27.5 | 27.5 | 27.5 | 27.5 | 27.5 | 27.5 | 27.5 |
| TrpRS | 27.5 | 27.5 | 27.5 | 27.5 | 27.5 | 27.5 | 27.5 |
| TyrRS | 27.5 | 27.5 | 27.5 | 27.5 | 27.5 | 27.5 | 27.5 |
| ValRS | 27.5 | 27.5 | 27.5 | 27.5 | 27.5 | 27.5 | 27.5 |
| IF1 | 27.5 | 27.5 | 27.5 | 27.5 | 27.5 | 27.5 | 27.5 |
| IF2 | 27.5 | 27.5 | 27.5 | 27.5 | 27.5 | 27.5 | 27.5 |
| IF3 | 27.5 | 27.5 | 27.5 | 27.5 | 27.5 | 27.5 | 27.5 |
| EF-G | 27.5 | 27.5 | 27.5 | 27.5 | 27.5 | 27.5 | 27.5 |
| EF-Tu | 837.5 | 837.5 | 837.5 | 837.5 | 837.5 | 837.5 | 837.5 |
| EF-Ts | 27.5 | 27.5 | 27.5 | 27.5 | 27.5 | 27.5 | 27.5 |
| **RF1** | **27.5** | **0** | **27.5** | **0** | **0** | **0** | **0** |
| **RF2** | **27.5** | **27.5** | **0** | **13.75** | **6.875** | **3.4375** | **1.71875** |
| RF3 | 27.5 | 27.5 | 27.5 | 27.5 | 27.5 | 27.5 | 27.5 |
| RRF | 27.5 | 27.5 | 27.5 | 27.5 | 27.5 | 27.5 | 27.5 |
| MTF | 27.5 | 27.5 | 27.5 | 27.5 | 27.5 | 27.5 | 27.5 |
| ADK | 27.5 | 27.5 | 27.5 | 27.5 | 27.5 | 27.5 | 27.5 |

| | | | | | | | |
|---|---|---|---|---|---|---|---|
| Ppiase | 27.5 | 27.5 | 27.5 | 27.5 | 27.5 | 27.5 | 27.5 |
| CK | 27.5 | 27.5 | 27.5 | 27.5 | 27.5 | 27.5 | 27.5 |
| NDK | 27.5 | 27.5 | 27.5 | 27.5 | 27.5 | 27.5 | 27.5 |
| T7 | 27.5 | 27.5 | 27.5 | 27.5 | 27.5 | 27.5 | 27.5 |
| **M.Alvus pyrrolysyl aaRS or M. Jannischii aaRS** | **0** | **27.5** | **27.5** | **27.5** | **27.5** | **27.5** | **27.5** |
| **Total Volume** | **1800** | **1800** | **1800** | **1786.25** | **1779.375** | **1775.938** | **1774.2188** |

**Table S2:** Buffer recipe for One Pot protein factor purification

| **Compound** | **Buffer A (mM)** | **Buffer B (mM)** | **Buffer HT (mM)** | **Stock Buffer A (mM)** | **Stock Buffer B (mM)** |
|---|---|---|---|---|---|
| HEPES (pH= 7.6) | 50 | 50 | 50 | 50 | 50 |
| Ammonium Chloride | 1000 | | | | |
| Magnesium Chloride | 10 | 10 | 10 | 10 | 10 |
| Potassium Chloride | | 100 | 100 | 100 | 100 |
| Imidazole (pH =7.6) | | 500 | | | |
| Glycerol | | | | 30% | 60% |
| B-mercaptoethanol | 7 | 7 | 7 | 7 | 7 |

**Table S3:** Buffer recipe for ribosome purification

| **Compound** | **Suspension Buffer (mM)** | **Suspension high salt buffer (mM)** | **Buffer C (mM)** | **Buffer D (mM)** | **Cushion buffer (mM)** | **Ribosome buffer (mM)** |
|---|---|---|---|---|---|---|
| HEPES (pH= 7.6) | 10 | 10 | 20 | 20 | 20 | 20 |
| Ammonium Sulfate (pH= 7.6) | | 3000 | 1500 | | | |
| Ammonium Chloride | | | | | 30 | |
| Magnesium Acetate | 10 | | 10 | 10 | 10 | 6 |
| Potassium Chloride | 50 | 100 | | | | 30 |
| Sucrose | | | | | 30 g/mL | |
| B-mercaptoethanol | 7 | 7 | 7 | 7 | 7 | 7 |

**Table S4:** 4x Energy mixture recipe

| Substance | Concentration |
|---|---|
| tRNA | 208 A260/mL |
| Amino acids | 1.2 mM |
| DTT | 4 mM |
| Potassium Glutamate | 400 mM |
| Spermidine | 8 mM |
| Creatine Phosphate | 80 mM |
| Folinic acid | 0.08 mM |
| ATP | 8 mM |
| GTP | 8 mM |
| UTP | 4 mM |
| Mg(OAc)2 | 47.2 mM |
| CTP | 4 mM |
| HEPES | 200 mM |

**Table S5:** Primers sequence to generate DNA template for tRNA and short peptide containing C-terminal FLAG epitote

| Name | Sequence |
|---|---|
| T7-F2 | GGCGTAATACGACTCACTATAG |
| F1-MjtRNA | TAATACGACTCACTATACCGGCGGTAGTTCAGCAGGGCAGAACGGC |
| R1-MjtRNA | CATGCGGATTTAGAGTCCGCCGTTCTGCCCTGCTGA |
| F2-MjtRNA | TGGTCCGGCGGAGGGGATTTGAACCCCT |
| fMKKKXAmber- f1 | TAATACGACTCACTATAGGGTTAACTTTAACAAGGAGAAAAACATG |
| fMKKKXAmber- r1 | GTCGTCGTCCTTGTAGTCCTACTTCTTCTTCATGTTTTTCTCCTTG |
| fMKKKX-r2-UGA termination codon | CGAAGCTCACTTGTCGTCGTCGTCCTTGTAGTC |
| fMKKKOchre- r1 | GTCGTCGTCCTTGTAGTCTTACTTCTTCTTCATGTTTTTCTCCTTG |
| fMKKK-r2-UAA termination codon | CGAAGCTTACTTGTCGTCGTCGTCCTTGTAGTC |
| fMKKK- r1-fixed | GTCGTCGTCCTTGTAGTCTCACTTCTTCTTCATGTTTTTCTCCTTG |

| | |
|---|---|
| MatRNAPyl8 -f1-ochre anticodon | GTAATACGACTCACTATAGGGGGACGGTCCGGCGACCAGCGGGTCT TTAAAACCT |
| MatRNAPyl8 -r1-ochre anticodon | TGGCGAGAGACCGGGGTGTCGAACCCCGCAAGGCTAGGTTTTAAA GACCCGCTGGT |
| MatRNAPyl8 -r2 | TGGCGAGAGACCGGG |
| MatRNAPyl6 -r1-opal anticodon | TGGCGAGAGACCGGGGCGTCGAACCCCGCTATGCTAGGTTTTGAA GACCCGCTGGTC |
| MatRNAPyl6 -f1-opal anticodon | GTAATACGACTCACTATAGGGGGACGGTCCGGCGACCAGCGGGTCT TCAAAACCT |
| Mb orche tRNA-f1-opal anticodon | GTAATACGACTCACTATAGGGAAACCTGATCATGTAGATCGAATGGA CTTCAAATC |
| Mb orche tRNA-r1-opal anticodon | TGGCGGAAACCCCGGGAATCTAACCCGGCTGAACGGATTTGAAGT CCATTCGATC |
| Mb orche tRNA-r2 | TGGCGGAAACCCCGGGAATCTA |

**Reporter sequences:**

Flag-peptide: X=amber or opal or ochre codon.

MKKKXDYKDDDDK

Super folded Green Fluorescent Protein (sfGFP):

MSKGEELFTGVVPILVELDGDVNGHKFSVRGEGEGDATIGKLTLKFICTTGKLPVPWPTL

VTTLTYGVQCFSRYPDHMKRHDFFKSAMPEGYVQERTISFKDDGKYKTRAVVKFEGDT

LVNRIELKGTDFKEDGNILGHKLEYNFNSHNVYITADKQKNGIKANFTVRHNVEDGSVQ

LADHYQQNTPIGDGPVLLPDNHYLSTQTVLSKDPNEKGTRDHMVLHEYVNAAGITWSH

PQFEKDYKDDDDK.

Super folded Green Fluorescent Protein containing in-frame ochre stop codon (X) (sfGFP):

MSKGEELFTGVVPILVELDGDVNGHKFSVRGEGEGDATIGKLTLKFICTTGKLPVPWPTL

VTTLTYGVQCFSRYPDHMKRHDFFKSAMPEGYVQERTISFKDDGKYKTRAVVKFEGDT

LVNRIELKGTDFKEDGNILGHKLEYNFNSHXVYITADKQKNGIKANFTVRHNVEDGSVQ

LADHYQQNTPIGDGPVLLPDNHYLSTQTVLSKDPNEKGTRDHMVLHEYVNAAGITWSH

PQFEKDYKDDDDK.

Orange fluorescent protein (OFP): X= amber codon

MSKGEELFTGVVPILVELDGDVNGHKFSVSGEGEGDATYGKLTLKFICTTGKLPVPWPTL

VTTLAXGLQCFARYPDHMKQHDFFKSAMPEGYVQERTIFFEDDGYYKTRAEVKFEGDT

LVNRIVLKGIDFKEDGNILGHKLEYNYNSHNVYIMADKQKNGIKVNFKIRHNIEDGSVQ

LADHYQQNTPIGDGPVLLPDNHYLSTQCALSKDPNEKRDHMVLLEFVTAAGITHDMDE

LYKGSGHHHHHH

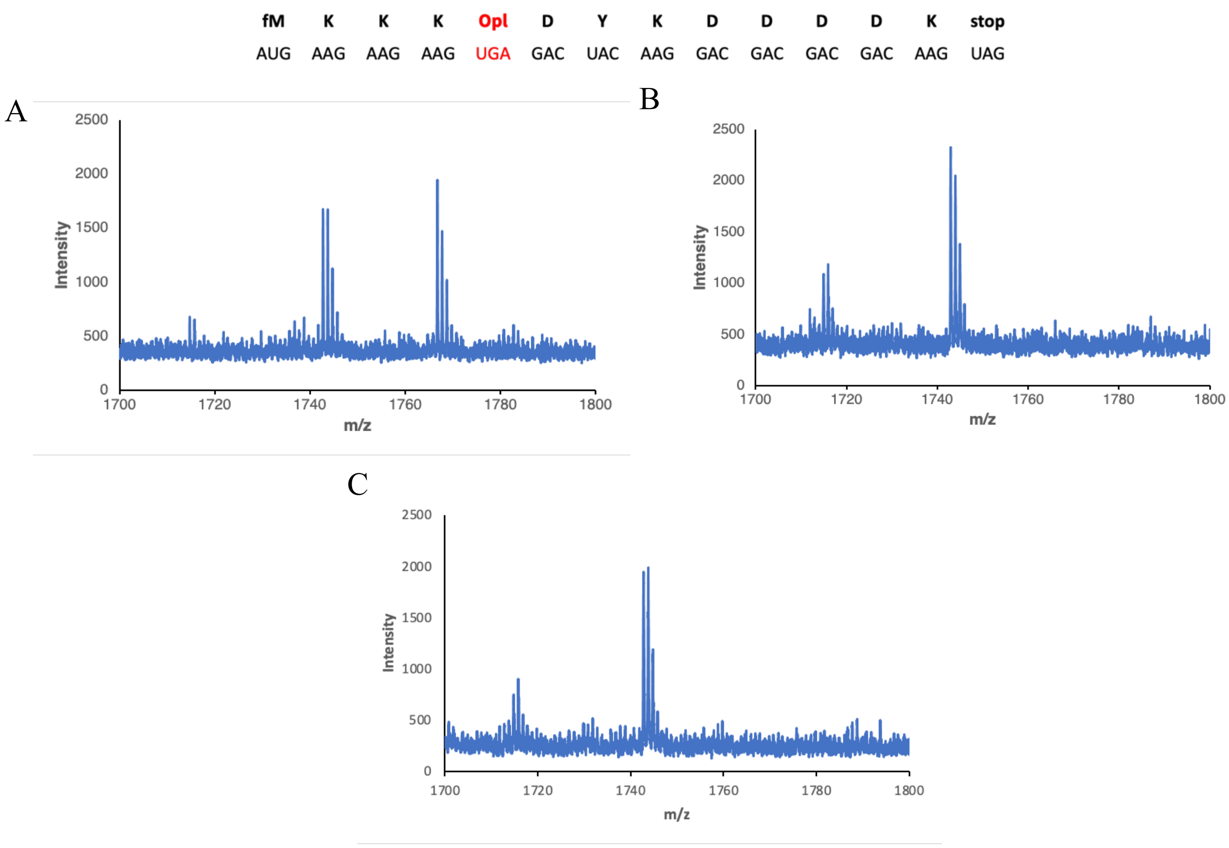


**Figure S6:** MALDI-MS report for opal codon (UGA) suppression to incorporate ProgK (A), and background (B- no ProgK) (C- no ProgK and no o-tRNA). Theoretical [$M+H^+$] for Trp misincorporation and ProgK are respectively 1742.81 m/z and 1766.83 m/z with the peptide reporter as indicated.

| fM | K | K | K | Ocr | D | Y | K | D | D | D | D | K | stop |
|---|---|---|---|---|---|---|---|---|---|---|---|---|---|
| AUG | AAG | AAG | AAG | UAA | GAC | UAC | AAG | GAC | GAC | GAC | GAC | AAG | UGA |

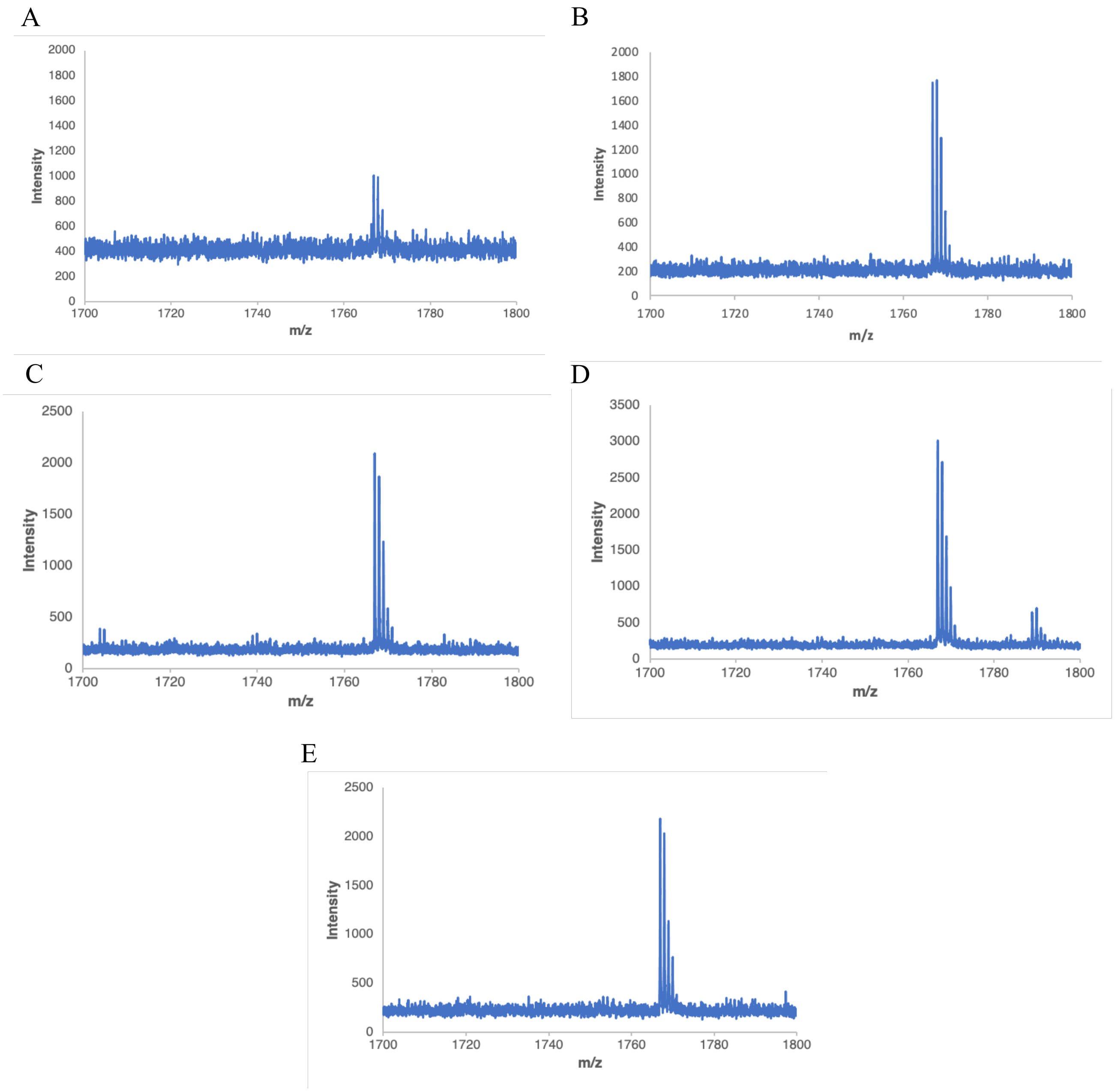


**Figure S7:** MALDI-MS report for ochre codon (UAA) suppression to incorporate ProgK (A- 1/1 RF2; B- ½ RF2; C- 1/4 RF2; D- 1/8 RF2; E- 1/16 RF2). Theoretical [M+H$^+$] for ProgK is 1766.83 m/z.
